# Pulled Diversification Rates, Lineages-Through-Time Plots and Modern Macroevolutionary Modelling

**DOI:** 10.1101/2021.01.04.424672

**Authors:** Andrew J. Helmstetter, Sylvain Glemin, Jos Käfer, Rosana Zenil-Ferguson, Hervé Sauquet, Hugo de Boer, Léo-Paul M. J. Dagallier, Nathan Mazet, Eliette L. Reboud, Thomas L. P. Couvreur, Fabien L. Condamine

## Abstract

Estimating time-dependent rates of speciation and extinction from dated phylogenetic trees of extant species (timetrees), and determining how and why they vary, is key to understanding how ecological and evolutionary processes shape biodiversity. Due to an increasing availability of phylogenetic trees, a growing number of process-based methods relying on the birth-death model have been developed in the last decade to address a variety of questions in macroevolution. However, this methodological progress has regularly been criticised such that one may wonder how reliable the estimations of speciation and extinction rates are. In particular, using lineages-through-time (LTT) plots, a recent study (Louca and Pennell, 2020) has shown that there are an infinite number of equally likely diversification scenarios that can generate any timetree. This has lead to questioning whether or not diversification rates should be estimated at all. Here we summarize, clarify, and highlight technical considerations on recent findings regarding the capacity of models to disentangle diversification histories. Using simulations we demonstrate the characteristics of newly-proposed “pulled rates” and their utility. We recognize that the recent findings are a step forward in understanding the behavior of macroevolutionary modelling, but they in no way suggest we should abandon diversification modelling altogether. On the contrary, the study of macroevolution using phylogenetic trees has never been more exciting and promising than today. We still face important limitations in regard to data availability and methodological shortcomings, but by acknowledging them we can better target our joint efforts as a scientific community.

## Introduction

A major goal in evolutionary biology is to understand the large-scale processes that have shaped biodiversity patterns through time. One important way to investigate this is by modelling species diversification using speciation and extinction, which can vary over time and among groups. It is commonplace to find areas, or clades, in phylogenetic trees that accumulate lineages faster than others. Diversification models often aim to explain this variation in diversification patterns by associating bursts of speciation or extinction with factors such as time (Höhna et al., 2016b), lineages (Rabosky, 2014), character traits (Maddison et al., 2007), or the environment (Condamine et al., 2013).

The growing number of large phylogenetic trees that capture a significant proportion of living species provide increasing power and resolution for such studies (Jetz et al., 2012; Smith and Brown, 2018; Upham et al., 2019). Furthermore, the availability of a wide variety of methods and software (e.g. BAMM (Rabosky, 2014), state-speciation and extinction (SSE) models (Maddison et al., 2007), RPANDA (Morlon et al., 2016), MEDUSA (Alfaro et al., 2009)) have made diversification studies increasingly popular in the last decade. Approaches that can link diversification to a particular process or trait are among the most appealing to researchers in the field because they enable us to test long-standing hypotheses in evolutionary biology and ecology. Examples include those related to the evolution of key innovations (Silvestro et al., 2014), the colonisation of new areas (McGuire et al., 2014), the effect of elevation (Lagomarsino et al., 2016; Quintero and Jetz, 2018) and the latitudinal diversity gradient (Rolland et al., 2014; Pulido-Santacruz and Weir, 2016; Rabosky et al., 2018; Igea and Tanentzap, 2020).

A recent study ((Louca and Pennell, 2020) abbreviated to LP) demonstrates how one approach, based on lineages-through-time (LTT) plots, cannot reliably estimate rates of speciation and extinction over time using extant timetrees. LP show how results of this approach can be misleading and provide potential solutions to the issues raised by proposing new summary statistics. This publication has already provoked a response from the community (e.g. Morlon et al. (2020)) and stimulated considerable discussion, with some going so far as to suggest that speciation and extinction rates cannot be estimated using phylogenetic trees (Pagel, 2020). As a result, this study has called into question the meaning of diversification rate estimates generated from any analytical framework. Here, we aim to outline the major concepts discussed in LP in an accessible way, targeting a broad audience. We then put the results and conclusions of LP into historical context and explore how the implications of this study apply to macroevolutionary modelling today.

## Modelling diversification rates

A typical workflow for diversification rate modelling using molecular phylogenetic trees is as follows. DNA sequence data are obtained for species in a study group, which are then used to estimate species relationships in the form of a phylogenetic tree. Typically, this phylogenetic tree contains only extant species, and it is time-calibrated using ages derived from different sources including fossils (Sauquet, 2013; Ho and Phillips, 2009). The output of this process is referred to as an extant timetree. Once a tree has been generated, a birth-death model is fitted to explain patterns of diversification in the tree. Note, however, that fossils are usually used for node calibration and tree shape estimation but are rarely incorporated in subsequent estimation of diversification rates, although recent methodological progresses now allow incorporating fossils as tips in the phylogenies (Ronquist et al., 2012; Heath et al., 2014) and birth-death models allow estimating rates of diversification Mitchell et al. (2019).

The simplest birth-death models assume that each branch of a phylogenetic tree shares the same rate of “birth” (speciation) events, as well as “death” (extinction) events (Nee et al., 1994; Nee, 2006; Ricklefs, 2007; Morlon et al., 2011). There are two principal parameters in the birth-death model, the speciation rate (*λ*) - the rate at which lineages arise, and the extinction rate (*μ*) - the rate at which lineages disappear. Under this simple framework *λ* and *μ* are constant over time (time-independent) and the same across all clades (clade-homogeneous). In addition, it is common that all extant taxa are not included in the phylogenetic tree, and the percentage of lineages present is known as the sampling fraction (or *ρ*) - the ratio of sampled species over the total species diversity for a given clade. By making use of these parameters, a birth-death model allows us to investigate whether the net diversification rate, defined as *r* = *λ − μ*, has varied over time or among clades (Morlon et al., 2011; Rabosky, 2014; Maliet et al., 2019; Barido-Sottani et al., 2020) and ultimately uncover the processes that have given rise to extant biodiversity in the study group.

## A summary of the main concepts and findings in Louca and Pennell (2020)

### The deterministic Lineages-Through-Time plot

The approach to study diversification used by LP relies on the Lineages-Through-Time (LTT) plot (Nee et al., 1992) (Fig. 1), which shows how extant lineages (i.e. only those existing in the present-day) accumulated over time using a phylogenetic tree. Each point in an LTT corresponds to a change in the number of lineages from the root of a phylogenetic tree at *t* = 0 to the present day at *t* = *T* (Fig. 1a). Alternatively, as in LP, time can be considered as an age (*τ* = *T − t*), where *τ* = 0 at the present and *τ* = *T* at the origin of the clade, or the root age (Fig. 1b). For consistency with LP, we will generally consider timescale as age (*τ*) in the equations we use throughout this manuscript.

**Figure 1.**
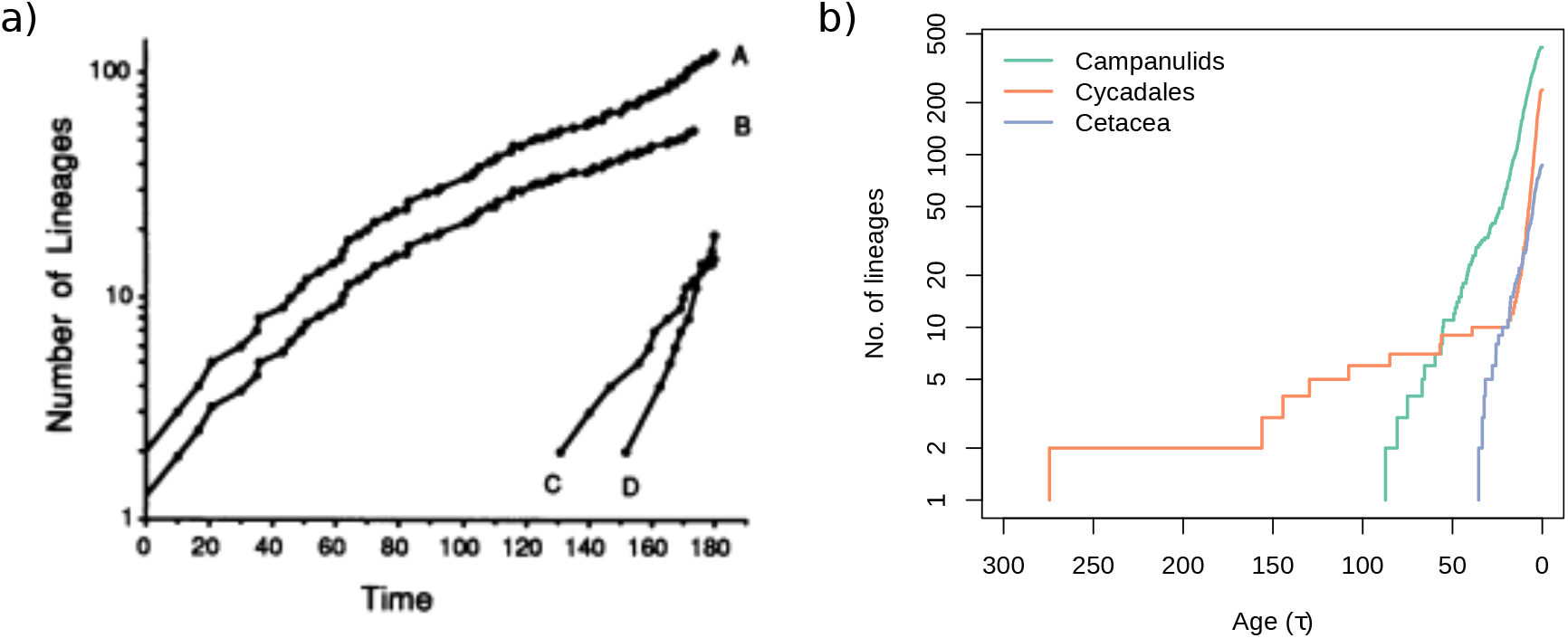
(a) The first example of a lineages-through-time plot (LTT), taken from Nee et al. (1992) and based on a phylogenetic tree of birds. On the y-axis is the number of lineages (log scale) and the x-axis is time since origin (present on the right hand side of the graph). *“Each point corresponds to a change in the number of lineages. Line A, the pattern of origination of all 122 lineages; line B, same as A, but without the Passeri (line C) and the Ciconliformes (line D). Line B has been shifted downward to aid visual comparison. The diversification rate is quantified by the steepness of the slope.”* In this panel, time is displayed from past to present as time since origin (*t*). (b) Three LTTs from modern phylogenetic trees of Campanulids (Beaulieu and Donoghue, 2013), Cycadales (Condamine et al., 2015) and Cetacea (Slater et al., 2010). In this panel time is shown from present to past as an age (*τ*).

Simply put, when a clade diversifies faster, the slope of the LTT becomes steeper, but when diversification slows, the slope of the LTT levels off. When only extant lineages are considered, as in LP, LTT plots will never exhibit a drop in total lineage diversity over time. Regardless of whether time is *τ* or *t* time in the equations, the LTT is usually plotted with the present on the right, thus its slope will never be negative. However, this does not mean that extinction does not have an effect on the shape of the LTT (Nee, 2006). By examining the shape of the LTT plot we can begin to understand how diversification rates fluctuated over the history of a clade (Ricklefs, 2007) and develop evolutionary hypotheses on why these fluctuations occurred.

To study general properties of phylogenetic trees, a model of the branching process is used. Several models are available, but the birth-death model is the most widely used, and is easily interpreted (Nee, 2006). The birth-death model is a continuous-time Markov chain where at any given age (*τ*) we can calculate the probability of speciation (birth of a lineage) or extinction (death of a lineage) happening. Because the birth-death process is stochastic, each run (i.e. simulation) will result in a different history of diversification, even if the probabilities for speciation and extinction are the same.

For such models we can calculate their expected value, either by averaging over multiple simulations or by approximating it with a set of continuous equations, yielding a deterministic model. Such a model produces the expected value one would get by averaging over an infinite number of simulations, thus it is deterministic because it is fully defined by the parameters, that is, no uncertainty from stochasticity is involved. This latter approach is widely used, and also taken by LP who model the birth-death process as a set of differential equations, which is advantageous because these equations can be solved analytically.

LP refer to an LTT generated by such models as a deterministic LTT (dLTT), which corresponds here to the expected LTT generated by trees with given speciation and extinction rates. Empirical LTTs generated using extant timetrees can be compared to model-generated dLTTs (where *λ* and *μ* are known) to disentangle how speciation and extinction have influenced patterns of diversity over time. To do this, the probability of the data given the model, or the likelihood, is calculated. Importantly, LP showed that, when *λ* and *μ* are clade-homogeneous across the tree, the likelihood can be fully written as a function of the observed LTT and the dLTT (see also Lambert and Stadler (2013)). Typically, by changing the parameters in the model, its dLTT resembles the empirical LTT to a greater or lesser extant, and the model is more or less likely. The best-fitting model can then be selected, representing our best hypothesis for how and to what extent speciation and extinction rates varied over time.

### Model congruence and congruence classes

Consider a simple model where *λ* and *μ* are fixed over time and among clades and all lineages have been sampled (*ρ* = 1). In this case, the slope of the LTT plot is *r* = *λ − μ*, except at times close to the present, where the effect of extinction diminishes and the slope becomes *λ* (Nee et al. (1994); see also Fig. 4 in Nee (2006)). If we know *λ* we can estimate *μ* by first estimating the slope of the LTT prior to the upward bend, which corresponds to *r* = *λ − μ*. This can then be rearranged to *μ* = *λ − r* and *μ* calculated using the estimated values of *λ* and *r* (in practice, both parameters can be inferred at once within a likelihood framework using equations in Nee et al. (1994)). LP develop upon this classical knowledge to show that if rates vary over time (*τ*) it is no longer possible to estimate *λ*(*τ*), as the value of *λ*(*τ*) at present does not yield any information about its past dynamic. In other words, it is possible to choose almost any historical scenario for *λ*(*τ*) and obtain a complementary scenario of *μ*(*τ*) that produces the same dLTT. If different models produce the same dLTT then they will also share the same likelihood for any given LTT.

**Figure 4.**
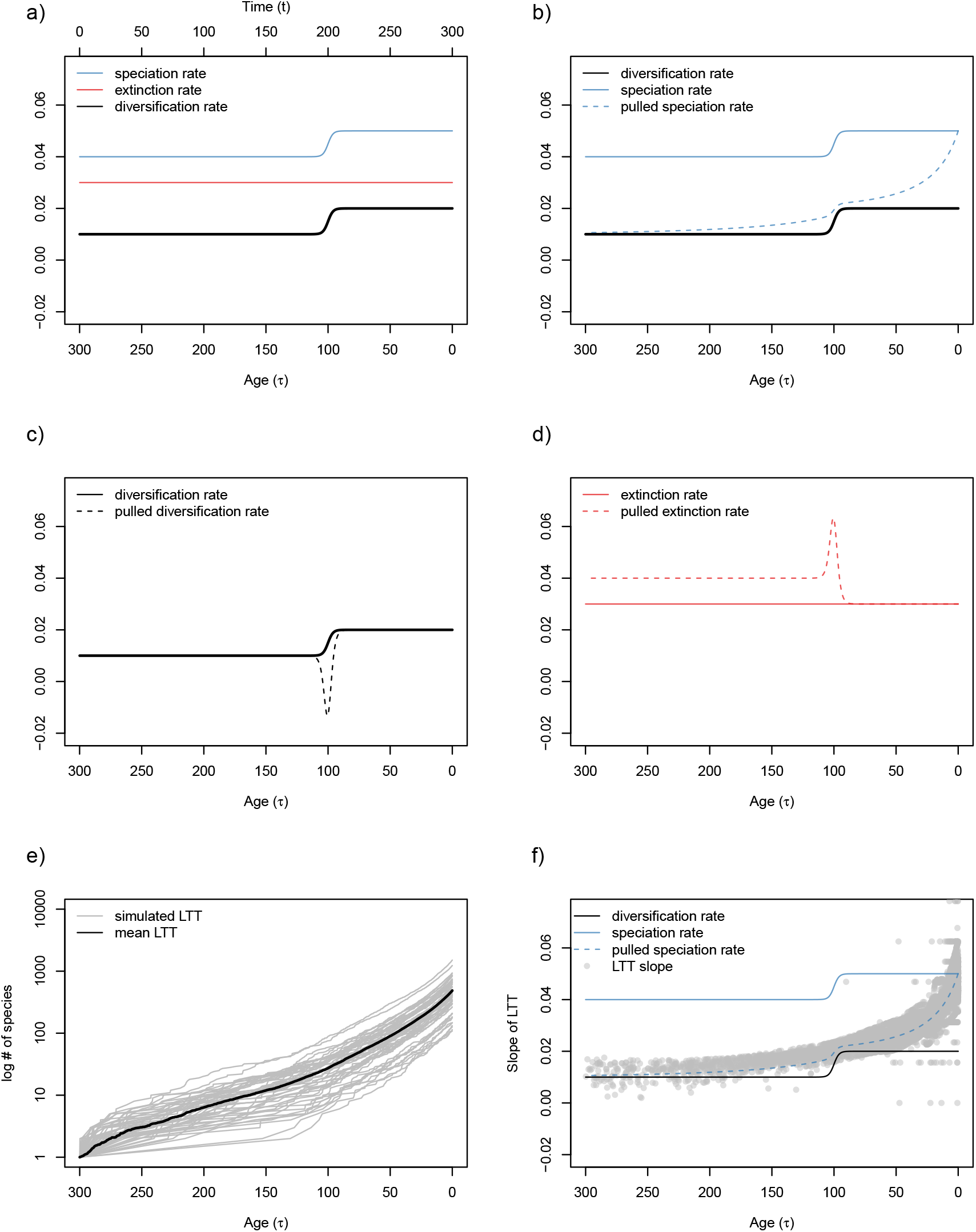
A slightly more complex example of the relationship between diversification rates and corresponding pulled rates where a single shift - an increase in speciation rate - has taken place. Panel (a) shows values of speciation rate (*λ*), extinction rate (*μ*) and diversification rate (*r*) over time. An additional axis, at the top of panel (a) shows time since origin (t). Panel (b) shows the gradual change in pulled speciation rate (*λ_p_*) during the shift in *λ*. Panel (c) compares *r* and pulled diversification rate (*r_p_*). The sudden increase in *λ* causes *r_p_* to decrease suddenly before recovering to the *r*. Panel (d) compares *μ* and pulled extinction rate (*μ_p_*) and shows an inverse pattern to panel (c). Panel (e) shows 50 LTT plots (grey lines) simulated with the parameters used in panels (a-d) and the mean LTT (black line). Panel (f) shows the slopes of the LTTs in panel (e) over time, matching *λ_p_* and again depicting the expected increase towards the present caused by the lack of effect of extinction.

LP call the set of models that generate the same dLTT a “congruence class”. These congruence classes contain an infinite number of models with different parameter values that all produce the same dLTT. LP explain that when trying to select the best model we often start with a relatively small set of allowed models that we test. For example, a set of two models where speciation rate is fixed and extinction rate is allowed to vary over time, or *vice versa*. This would produce two equally likely models when trying to explain a slowdown in diversification, one indicating the case was an increase in extinction rate, the other a drop in the speciation rate - there is no way of distinguishing between them Crisp and Cook (2009); Burin et al. (2019). LP suggest that instead of selecting the model closest to the true process, we are instead selecting the model closest to the congruence class that includes the true process (see Fig. 3 in LP). In extreme cases, the best fitting model could thus be further from the true process than a more correct model, just because the former is included in the congruence class and the latter is not. However, LP concede that because we only assess a limited set of models, it is unlikely that we encounter models belonging to the same congruence class, but it is nevertheless possible. The consequence of multiple, equally likely models with different speciation and extinction rates is that these rates cannot be determined. This is a statistical phenomenon known as unidentifiability - the likelihood is the same for multiple parameter values making it impossible to choose one over another.

**Figure 3.**
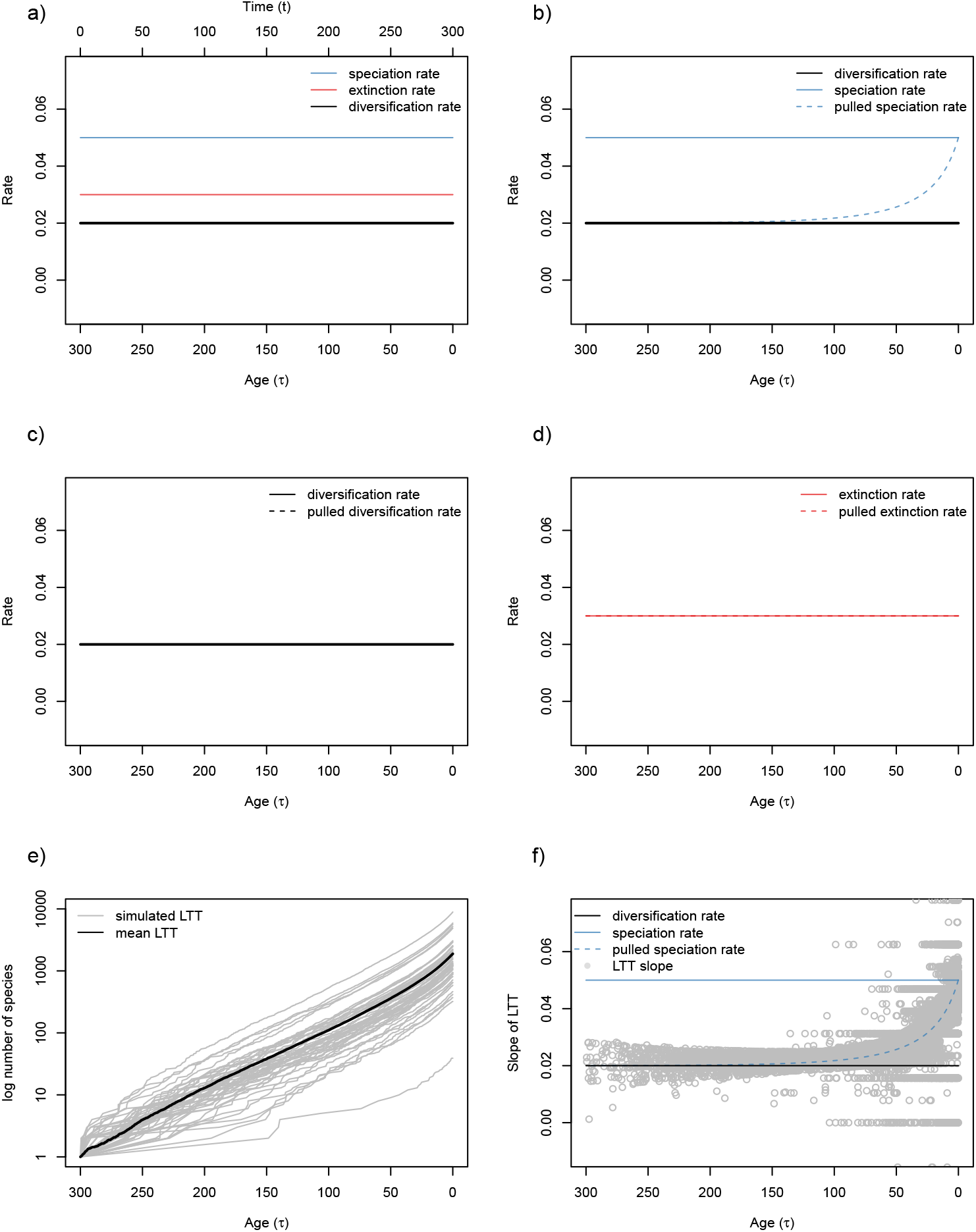
A simple example of the relationship between constant diversification rates and corresponding pulled rates. Panel (a) shows values of speciation rate (*λ*), extinction rate (*μ*) and diversification rate (*r*) over time. An additional axis, at the top of panel (a) shows time since origin (t). Panel (b) shows how in the past, pulled speciation rate (*λ_p_*) is identical to the diversification rate (if sampling fraction = 100%) while closer to the present *λ_p_* approaches speciation rate. The following two panels compare (c) *r* & pulled diversification rate (*r_p_*) and (d) compares *μ* & pulled extinction rate (*μ_p_*). In these two cases the pulled rates are identical to the traditional rates. Panel (e) shows 50 LTT plots (grey lines) simulated with the parameters used in panels (a-d) and the mean LTT (black line). Panel (f) shows the slopes of the LTTs in panel (e) over time, matching *λ_p_* and depicting the expected increase towards the present caused by the lack of effect of extinction - lineages do not have enough time to go extinct towards the present. An interactive version of this plot, in which parameters can be modified, can be found at https://ajhelmstetter.shinyapps.io/pulledrates/.

### Unidentifiability

In macroevolutionary modelling we might be interested to know how both *λ* and *μ* have changed over time (Alfaro et al., 2009). The unidentifiability issue outlined above means that we would not be able to ascertain the true parameter values of the models that generate our dLTTs. Another well-known example of this in macroevolution is the unidentifiability of *α* and *θ* from Ornstein-Uhlenbeck models of trait evolution (Ho and Ané, 2014). This problem is not unique to macroevolutionary models, and, in fact, stems from a basic mathematical issue (Rannala, 2002; Ponciano et al., 2012).

Consider a simple example of the concept in which we want to determine the parameter values for *x* and *y*. For each value of *x* in equation 0.1 below, we can find a *y* that satisfies this equation - and there are an infinite number of equally likely possibilities. It is only when we add more information (in the form of equation 0.2) that we can determine the unique pair of values for *x* and *y*. Put simply, a solution can be found only if you possess at least the same number of equations as unknowns. In this case the unidentifiability is caused by overparameterization - there is an excess of parameters such that the model cannot estimate the values of any of them.

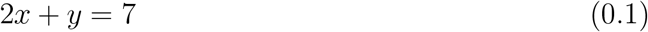

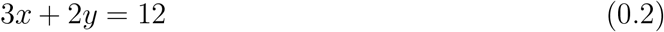

Though the LTT is generated through the use of many different observations and elements (DNA, fossils for time-calibration, extant species sampling) it is represented by a single curve made up of one observation at any given point in time that represents the number of lineages in a clade (Fig. 1). Fitting a model to an LTT is like fitting two parameters (*a* and *b*) for the slope (*a − b*), which gives you only one value. If we try to estimate *a* and *b* separately we find it impossible (Fig. 2a,b). However, we can estimate *a − b* (Fig. 2c). Estimates of *a* and *b* are fully correlated (Fig. 2d) and we find a flat surface in the likelihood where different pairs of values for *a* and *b* are equally likely (Fig. 2e), signifying unidentifiability.

**Figure 2.**
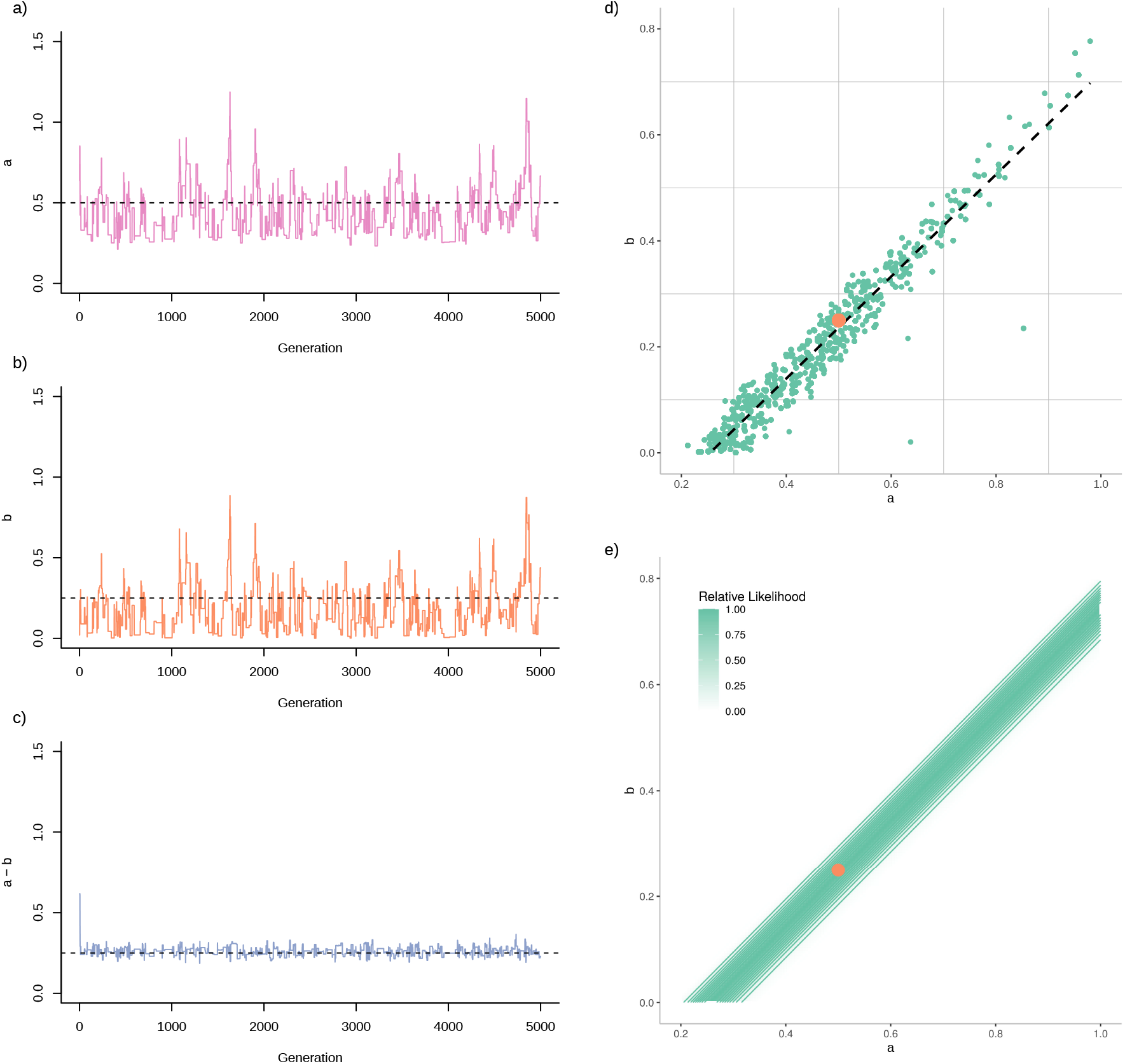
An example of unidentifiability issues encountered when trying to estimate values of two parameters (*a* & *b*) for the slope *a − b*. We used a Bayesian Monte-Carlo Markov Chain approach to try to estimate the values of *a* and *b* separately. We ran a chain for 5000 generations, sampling each generation. The traces for (a) *a* and *b* show a great deal of uncertainty in the parameter estimates compared to the estimates for (c) *a − b*. True values are shown as black dashed lines in panels (a-c) and as orange circles in panels (d-e). We plotted *a* against *b* and found that they two parameters were highly correlated (d). When then calculated the relative likelihood over a range of parameters values and found a flat ridge in the likelihood where different pairs of values for *a* & *b* are equally likely - or unidentifiable (e).

This problem has been highlighted previously (Nee, 2006), where *a − b* is the net diversification rate (*r* = *λ − μ*). However, the birth-death model is more complex than the example illustrated in figure 2. As explained above, speciation and extinction rates are actually identifiable when time-independent because the slope of the LTT reaches *λ* at the present. Our ability to reliably estimate these traditional diversification rates (*λ*, *μ* & *r*) depends on the amount of information we have available, and the assumptions we make in our model. For example, if the sampling fraction (*ρ*) is not known (or assumed) we can no longer reliably estimate *λ* and *μ* because this third unknown parameter in the model leads to unidentifiability (Table 1). However, *r* = *λ − μ* (with *ρ*) remains identifiable, as the system is reduced to two parameters only. Likewise, as LP show, if we relax the assumption of constant rates and allow *λ* and *μ* to vary over time, then all traditional parameters become unidentifiable, including *r*(*t*), even if *ρ* is known or assumed. (Table 1).

**Table 1.** A table detailing the parameters we can estimate with the Lineages-through-time plot (LTT) approach used in Louca and Pennell (2020) when rates are either constant or time-dependent. When speciation and extinction rate are constant we are able to infer all traditional (*r*, *λ*, *μ*) and pulled rates (*rp*, *λp*, *μp*). If sampling fraction (*ρ*) is unknown, we lose the ability to infer *λ* and *μ*. If *λ* and *μ* vary over time only pulled rates remain identifiable.

|  | Speciation | Extinction | Sampling fraction | Ref | Identifiable parameters |
| --- | --- | --- | --- | --- | --- |
| constant rates | $\lambda = a$ | $\mu = b$ | $\rho = 1$ | Nee et al. (1994) | $r, \lambda, \mu, r_p, \lambda_p, \mu_p$ |
| | $\lambda = a$ | $\mu = b$ | $\rho < 1$ (known) | Nee et al. (1994) | $r, r_p, \lambda_p, \mu_p$ |
| | $\lambda = a$ | $\mu = b$ | $\rho < 1$ (unknown) | Stadler (2013) | $r, r_p, \lambda_p, \mu_p$ |
| time-dependent rates | $\lambda = f(\tau)$ | $\mu = g(\tau)$ | $\rho = 1$ | Louca and Pennell (2020) | $r_p, \lambda_p, \mu_p$ |
| | $\lambda = f(\tau)$ | $\mu = g(\tau)$ | $\rho < 1$ (known) | Louca and Pennell (2020) | $r_p, \lambda_p, \mu_p$ |
| | $\lambda = f(\tau)$ | $\mu = g(\tau)$ | $\rho < 1$ (unknown) | Louca and Pennell (2020) | $r_p, \lambda_p, \mu_p$ |

To exemplify the problem, LP used a very large angiosperm phylogenetic tree (Smith and Brown, 2018) to show that the observed LTT is congruent with two opposing scenarios (Fig. 2 in LP): either a continuous increase or a continuous decline in both *λ*(*τ*) and *μ*(*τ*) (though the resulting diversification rates of these two scenarios are very similar). Therefore, if we observe a rapid increase in the number of lineages in our LTT (Fig. 3) we cannot determine if it was caused by a decrease in extinction rate, or an increase in speciation rate. If we want to use models to explain LTTs then traditional variables are inadequate and we must look towards other possible solutions.

### Pulled rates and their interpretation

LP’s solution is to use the approach described in Louca et al. (2018), namely not to estimate *λ*(*τ*), *μ*(*τ*) and *ρ*, but “pulled” rates that can be directly measured from the shape of the LTT. There are three pulled rates (*λ_p_*, *μ_p_*, *r_p_*) in Louca et al. (2018). These pulled rates are based directly on the dLTT - they make use of the slope at a given time (*τ*) and the change in the slope, or curvature of the plot. Thus, any dLTT yields a unique set of pulled rates that summarise a congruence class, thereby eliminating the unidentifiability issue. However, these rates are not the speciation and extinction rates everyone knows - so what are they and how are they different from traditional rates?

An important consequence of using extant timetrees when investigating patterns of diversification is that LTT plots will likely underestimate the number of lineages at any given time because our trees are missing species (Ricklefs, 2007; Silvestro et al., 2018). Species can be missing for two reasons: (1) they went extinct, or (2) they are extant but were not sampled. However, these two factors will have different effects on the LTT and our estimates of diversification rates. Extinction must have occurred in the past. Lineages that originated recently have had less time to go extinct (Nee et al., 1994; Ricklefs, 2007), so the effect of extinction on our estimates using only extant species is reduced towards the present. As mentioned above, this leads to an increase in the rate of lineage accumulation towards the present, as the effect of extinction decreases, which occurs even when rates are constant (Fig. 3). Conversely, incomplete sampling of a group occurs up to the present day and more strongly affects estimates of recent history (Phillimore and Price, 2008; Heath et al., 2008; Cusimano and Renner, 2010), as the deeper nodes in the phylogeny can be reconstructed with only a few species. The relative importance of extinction and sampling fraction will influence whether *λ_p_* departs from *λ* more in the past or in the present. To summarise, the presence of extinction will cause us to underestimate speciation rate further in the past, because the number of extinct species increases as we consider more time, while incomplete sampling will lead to underestimates of speciation rates that are more recent (Heath et al., 2008; Cusimano and Renner, 2010; Brock et al., 2011).

Formally, at a given time (*τ*), *λ_p_* is the estimated speciation rate multiplied by 1 minus the probability that a lineage is missing from the tree due to extinction or incomplete sampling, *E*. We will not go into details regarding the calculation of *E* here, but further information can be found in supplementary materials of LP. *λ _p_* is calculated using the following equation:

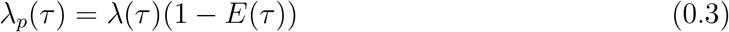

So, if all species are in the tree and there is no extinction (i.e. the probability of missing lineages, or *E*, is 0) then the *λ_p_* is equal to the (un-pulled) speciation rate, *λ*. Any increase in extinction rate or the number of unsampled lineages (i.e. *E >* 0) will cause *λ_p_* to drop, or be “pulled”, below speciation rate (Fig. 3). The lower the extinction rate and the greater the sampling fraction, the closer the estimate of *λ_p_* will be to *λ*.

LP also use pulled diversification rate (*r_p_*), a parameter that is similar to the net diversification rate (*r* = *λ − μ*) but is again modified by an other term. This new term is the relative 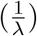 rate of change in speciation rate over time 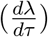. This causes the pulled diversification rate to lag behind the unpulled rate. The “pull” of *r_p_* is actually a delay in the response of this parameter when compared to diversification rate. This is in contrast to the “pull” of *λ _p_*, which refers to a reduction in the estimated value of *λ _p_* relative to *λ*. Pulled diversification rate can be represented by the following equation:

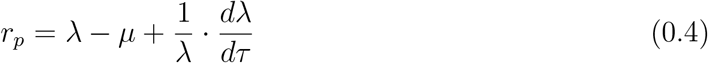

Consider an example where we have an increase in speciation rate at around 100 Ma in a clade (Fig. 4). When considering time as an age (using *τ*), speciation rate increases as *τ* decreases from the origin of the clade (*τ* = 300 Ma) to the present (*τ* = 0). This means that when speciation accelerates, *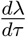* is negative. This leads to a “drop” in *r*_p_ *dτ* (Fig. 4c) before it stabilizes at a new value of of *r_p_* that is higher than the previous value, reflecting the increase in *λ*. However, the change in the slope of the LTT (Fig. 4e,f) is minimal, so this is not actually measurable from a phylogenetic tree. We note that LP also defined a pulled extinction rate, (*μ_p_*), which is similar to *r_p_* so we do not discuss it here (see LP, Louca et al. (2018) for further details).

The difference between the true diversification rate and an estimated *r_p_* can be likened to a race between an amateur and a professional race car driver. The professional driver, representing the true diversification rate in our analogy, hits the apex of each corner, going smoothly around a racetrack until the finish line. The amateur, representing *r_p_*, will eventually arrive at the finish line, but may exceed track limits a few times when doing so because of their poor reactions. However, if the track is simply a straight line both will perform equally well. This is because the *r_p_* is equal to the diversification rate (*r* = *λ − μ*) whenever *λ* is constant in time *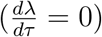*, but differs from *r* when *λ* varies with time (see Technical considerations below for more details).

With these new variables we can revisit questions such as: has diversification been constant over time? Pulled rates can be estimated with many commonly used models of diversification (Louca and Pennell, 2020). For example, *λ_p_* is the speciation rate one would get by constraining extinction to be 0 and assuming complete species sampling. For *r_p_* this involves estimating *r* by making *λ* time-independent. In summary, *λ_p_* provides information about how *λ* changes over time while taking into account past extinction and the proportion of lineages sampled. *r_p_* provides a slightly delayed estimate of *r* with extreme responses to rapid changes in *λ*. While *λ_p_* can be very different from the underlying speciation and extinction rates, *r_p_* is close to the net diversification rate as long as *λ* does not change too rapidly.

## Technical considerations

### How continuously can speciation and extinction rates vary?

Although this is standard practice, it should be noted that the approach of LP considers speciation and extinction to be continuous processes: at any infinitesimal time interval, the species number changes infinitesimally through speciation and extinction. In the birth-death process, however, the smallest amount of change in the number of species is one, and this happens only at particular moments in time. Even if speciation, in reality, is a complex process that takes time (Etienne and Rosindell, 2012), it is sufficiently fast to be considered instantaneous in evolutionary time. An empirical LTT plot will thus show discrete events, rather than being a continuous function, as is the dLTT. To measure the pulled rates, LP propose calculating the slope and curvature of the LTT plot. For the dLTT, where the number of lineages is a continuous function of time, these are the first and second derivative of this function. For empirical LTT plots, one has to calculate the slope and curvature using some time interval. When working with a large phylogenetic tree and many species (as in the examples discussed by LP and (Louca et al., 2018)), the LTT is smooth and the slope and curvature, which are necessary for the estimation of the pulled rates, can be reliably estimated. However, many studies attempt to estimate diversification rates with relatively small numbers of species (i.e. *<* 1000 (Hutter et al., 2017) or even *<* 100 Duan et al. (2018)). Thus, as the number of species diminishes, one has to be aware that overparameterization might occur, and it would be wiser to stick to simple functions of diversification (or their pulled variants) through time, the simplest being time-independent rates. Furthermore, even in large trees, rates will always be estimated using a time interval that contains a sufficient number of speciation and extinction events. The consequence of this is that rapid changes in diversification rates might be missed due to the resolution of the chosen interval.

### The “pull” in r_p_ is a result of the lag time between extinction and speciation

Consider a simple case with no extinction (*μ* = 0) so that changes in *r* that only come from changes in *λ*. If so, *r* = *λ* but *r_p_* is not exactly *λ* because of temporal variations in *λ* (the term 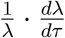 in equation (0.4)). LP suggest that “*the pulled diversification rate can be interpreted as the effective net diversification rate if λ was time-independent*”. In our example, this means replacing a scenario where *μ* is constant (at 0) and *λ* varies with a scenario where *λ* is constant and *μ* varies, as in LP. The difficulty of using *μ* to explain variation is that there is a slight delay between the effect of speciation and the effect of extinction. It is necessary to wait for species to arise before they can go extinct.

As mentioned previously, lineages that originated more recently have had less time to go extinct. In a constant birth-death process, this is only visible in recent history: the slope of the LTT is *r* = *λ − μ* during most of the past but increases to *λ* for very recent times where the stationary behaviour has not yet been reached (Fig. 3). However, this phenomenon is not unique to very recent times - it will also occur whenever there is a change in speciation rate. Ultimately, this is the cause of the difference between *r_p_* and *r*.

To illustrate this, imagine a massive increase in the number of lineages caused by a burst of speciation (Fig. 4). Over a short time period many new lineages have become available for potential extinction but they have yet to go extinct because not enough time has passed since they appeared for extinction to take place. There is now a disequilibrium between speciation and extinction, manifested as a lag in the time extinction takes to affect all of the new lineages. As time continues, these numerous new lineages will begin to go extinct, meaning that frequency of species extinction will increase to “catch up” to speciation and reach a new stationary point. This effect is stronger when *λ* varies rapidly (i.e. high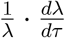). Conversely, speciation cannot occur in a lineage after it has gone extinct, so there is no similar lag caused by changes in extinction rate. This is also why variation in extinction rate would not cause *r_p_* to deviate from *r* (Fig. 5b).

**Figure 5.**
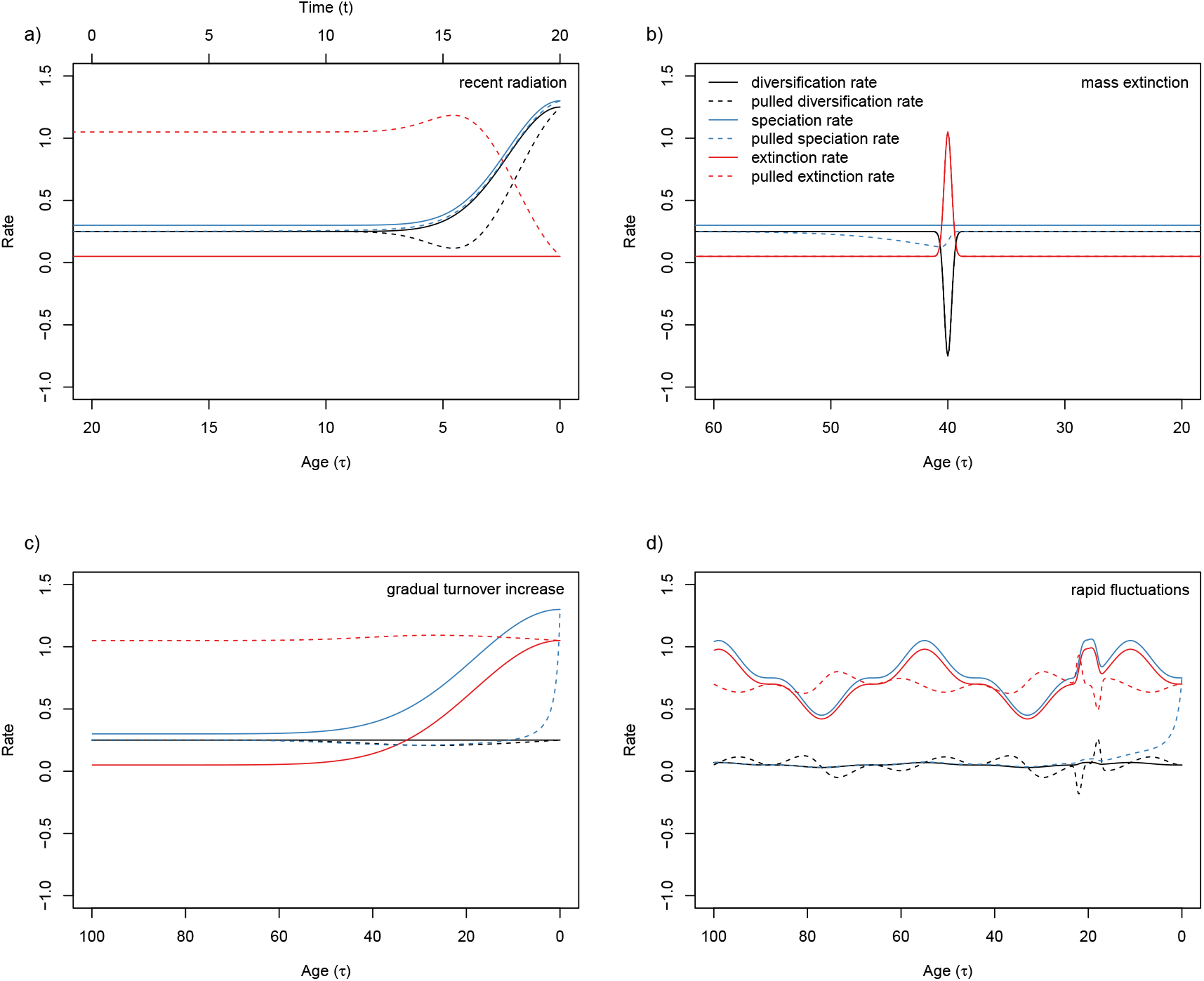
Comparison of traditional and pulled rates under three simulated diversification scenarios that are commonly investigated (a-c) and a final, more complex scenario. Panel (a) shows a recent radiation where diversification rate and speciation rate sharply increase towards the present. An additional axis, at the top of panel (a) shows time going forward (t). Panel (b) shows a mass extinction event at 40 Ma in which extinction rate briefly but rapidly increases and then falls back to previous levels. Panel (c) shows a gradual increase in species turnover rate (both speciation and extinction rate increase slowly over time). Panel (d) shows a scenario where speciation and extinction rates are similar to each other but are in rapid fluctuation over time. This results in a relatively constant diversification rate (*r*) and a rapidly fluctuating pulled diversification rate (*r_p_*) that remains close to *r*.

## Discussion

A recent study by Morlon et al. (2020) presents an alternative point of view that opposes the conclusions in LP. They focus on how a hypothesis-based framework allows us to overcome many of the issues that are raised in LP. Indeed, we are limiting our set of models to be tested to only those that represent our hypotheses about the factors shaping diversification in a given group. We are not often interested in determining the precise values of speciation and extinction rate but rather how different diversification scenarios summarised by models containing *λ* and *μ* explain patterns in a phylogenetic tree. The criticisms put forward by Morlon et al. (2020) will stimulate important discussion of considerations when using diversification models. We extend this discussion by highlighting several key points that must be considered in addition.

### Uses and limitations of LTTs

LTTs are a simplistic way to visualize and summarize a time-calibrated phylogenetic tree, ignoring the full distribution of branch lengths, tree topology and extinct species (Morlon et al., 2011). However, under the assumption that *λ* and *μ* are clade-homogeneous, LP showed that the LTT contains the complete information about the underlying branching process (See also Lambert and Stadler (2013)). This simplicity provided the opportunity for LP to show mathematically how LTTs can lead to misinterpretation. These issues are not new to macroevolutionary biology. A review by Nee (2006) clearly demonstrated how an LTT may change when extinction is present alongside speciation (birth-death), as opposed to speciation alone (pure-birth), summarising theory from previous works (Nee et al., 1992, 1994; Harvey et al., 1994). If the growth of an extant timetree is represented as an LTT on a semi-log scale (i.e. lineage number is logarithmic, time is not, see Fig. 1) we would expect the trend to be linear under a pure-birth process (with constant speciation and no extinction). If extinction is introduced then the LTT would deviate from this linearity. When both rates are constant and greater than 0, the curve is expected to be linear over most of its history, but as time reaches the present the rate of lineage accumulation will increase (i.e. the LTT slope will become steeper), as shown in Figure 3a. With no prior knowledge of the parameters, this could be, at least in part, because of increasing speciation rate towards the present (Fig. 3b), instead of decreasing effect of extinction (Fig. 3). It is important to keep in mind that we are dealing with a phylogenetic tree made up of entirely extant species. The unobserved branches of species that went extinct (and are therefore not in the extant timetree) do not contribute to the lineage counts in the LTT, making the estimated lineage accumulation rate lower in the past (or “pulling” it down). Nee et al. (1994) highlighted this issue 20 years ago in the context of models where diversification rates were constant over time and now LP have provided an important extension of this idea to models that allow for rates to vary through time. Since Nee et al. (1994), the well-known limitations of LTTs for inferring speciation and extinction rates have continued to be addressed in other studies (Ricklefs, 2007; Vamosi et al., 2018; Rabosky and Lovette, 2008; Crisp and Cook, 2009), and most recently in LP.

### Diversification rates vary among clades

The conclusions of LP imply that we can test hypotheses about whether diversification rates deviate from constancy over time using pulled rates. We would be unable to pin this on changes in speciation or extinction rates, but would get a sense of how variable diversification has been (Burin et al., 2019). This would be useful for testing whether diversification in particular clades has remained constant or been subject to large shifts in diversification (e.g. mass extinctions) but not when diversification rate has shifted in a subclade (e.g. due to the evolution of a key innovation). The first use of pulled rates was in Louca et al. (2018), where they studied bacterial diversification, stating “*Our findings suggest that, during the past 1 billion years, global bacterial speciation and extinction rates were not substantially affected during the mass extinction events seen in eukaryotic fossil records.*” This might suggest that nothing particularly extraordinary happened in the macroevolutionary dynamics of bacteria in the last billion years. However, it is important to note that the models used in Louca et al. (2018) and in Louca and Pennell (2020) do not allow rates to vary among clades. The rates estimated using such clade-homogeneous models will correspond to the average rates over time in the entire study group, therefore missing out on any variation among clades - for example any difference in diversification rates between those species that use terrestrial *versus* marine environments (Louca et al., 2018). Subclades are important in driving inferred diversification patterns (see Morlon et al. (2011); Maliet et al. (2019); Rabosky (2020)), so this may mean that we miss out on influential and interesting dynamics when using pulled rates. Louca et al. (2018) touch on this point themselves: “*It is possible that diversification within individual bacterial clades may have been influenced by eukaryotic radiations and extinctions, and that these cases are overshadowed when considering all bacteria together.*” Given the diversity of life on Earth, it is unrealistic to assume that major events would have had the same effect on all lineages of a large, cosmopolitan clade, with vast amounts of genetic, morphological and ecological variation. Such an assumption prevents us from investigating some of the most interesting and fundamental questions in macroevolutionary biology e.g. why are some clades more diverse than others?

The same criticisms could be levelled at LP’s use of a large phylogenetic tree of angiosperms (Smith and Brown, 2018) that contains more than 65,000 of the roughly 300,000 known species, ranging from small ephemeral plants like *Arabidopsis thaliana* to gigantic, long-lived trees such as *Eucalyptus regnans*. Furthermore, a large amount of research has shown that diversification rates have varied significantly among flowering-plant clades (e.g. O’Meara et al. (2016); Igea et al. (2017); Vamosi et al. (2018); Onstein (2019); Soltis et al. (2019); Zenil-Ferguson et al. (2019); Magallón et al. (2019)), and pulled rates would not be able to contribute to furthering our understanding of why this may be.

Fortunately, the assumption of homogeneous rates among clades is not common in modern approaches. For instance, Bayesian Analysis of Macroevolutionary Mixtures (BAMM) (Rabosky, 2014) is one of several methods (Alfaro et al., 2009; Morlon et al., 2011; Höhna et al., 2016a; Maliet et al., 2019; Barido-Sottani et al., 2020) that relaxes the assumption that all lineages share the same evolutionary rates at a given point in time (Rabosky, 2017). This is a key difference from the models used by LP because it allows lineages to differ in their rates of speciation and extinction. With BAMM, the entire phylogeny could be described using a model similar to what is used in Louca and Pennell (2020), or alternatively, it could be described using multiple processes that explain rates of diversification on different parts of the tree. Within each of these processes, *λ* and *μ* probably still faces the same unidentifiable issues outlined in LP. However, BAMM makes use of the full topology that includes information (e.g. branch lengths) that the LTT lacks. A model that allows diversification rate varies among clades could yield entirely different insights compared to a model where this rate can only vary in time.

Another model commonly used to estimate and compare diversification rates among clades is the Binary-State Speciation and Extinction (BiSSE) model (Maddison et al., 2007), part of a family of models known as the state-dependent models of diversification (-SSE models (Ng and Smith, 2014; O’Meara and Beaulieu, 2016; Beaulieu and O’Meara, 2016; Caetano et al., 2018)). These models are extensions of the birth-death model that also includes information about character states of extant species. SSE models jointly estimate ancestral states at each node of the phylogenetic tree, rates of transition between character states, and state-dependent diversification rates. LP state that the likelihood functions of SSE models are too complex to be addressed in their study, but suggest that the same problems they uncover probably still apply. As in BAMM, BiSSE makes use of the full tree topology (Maddison et al., 2007) - it also considers character state evolution, rather than just the timing of branching events as in the LTT (Nee et al., 1994). LP further suggest that it remains unclear how the dependence on character states (which, if removed, collapses equations in BiSSE to those shown in Nee et al. (1994)) affects the unidentifiability issue they raise. In the original BiSSE model (Maddison et al., 2007), two important and relevant assumptions were made: (1) sampling fraction is assumed to be 100%, and (2) speciation, extinction and transition rate remain constant through time per character state.

These may allow the BiSSE model to overcome (or pre-empt) some of the problems raised by LP. LP show that *λ* equals *λ_p_* when sampling fraction is 100% and *μ* = 0. The first of these was assumed in the original BiSSE model, though it has since been relaxed (FitzJohn et al., 2009). Extinction can easily be set to 0 in these models, which satisfies the second BiSSE assumption and allows estimation of *λ* (e.g. Joly and Schoen (2021)). Similarly, *r_p_* equals *r* when *λ* is constant, also an assumption in BiSSE. With these additional assumptions, the congruence class collapses to only one model. However, this does not mean that BiSSE actually estimates the speciation rate of a clade correctly; LP’s results show we should take this BiSSE estimate as a proxy for the diversification rate. Nevertheless, we stress that the likelihood of time-dependent diversification models (as in LP) is not the same as the likelihood of state-dependent diversification models (-SSE models) and what is unidentifiable in the former does not tell us about identifiability in the latter.

It is unclear how clade-heterogeneous rates would affect model congruence, and how the additional information included when using models included in programs such as BiSSE and BAMM would affect the unidentifiability issues. However, what is clearer is that the issues raised in LP cannot be readily applied to commonly used macroevolutionary approaches without further work to show that criticisms related to LTT-based approaches are applicable to these more complex models. Even if unidentifiability issues remain in such models they may not be relevant to the questions the models were built to answer, for example those models that test for variation in diversification rates in association with particular clades or traits. In cases like these, it is not the precise values of rates that are important but instead whether rates in one group of lineages are higher than another.

Perhaps most importantly, this means that we should not forego building models that estimate diversification rates because one, simplistic approach has problems known from a long time, but instead continue to improve them and build upon the work done in LP. A case in point is the issue of null model choice when using SSE models raised by Rabosky and Goldberg (2015). This criticism spurred on innovation that led to the development of models with hidden states (Beaulieu and O’Meara, 2016), which are now present in various new incarnations of the SSE approach (e.g. (Caetano et al., 2018; Herrera-Alsina et al., 2019)).

### Pulled rates are difficult to interpret

LP compared the usefulness of pulled rates to effective population size (*N_e_*) in population genetics. *N_e_* can be broadly defined as the number of breeding individuals in an idealised population (e.g. constant size, random mating etc.) that would be able to explain the summary statistics of an observed population (e.g. amount of polymorphism, level of inbreeding). *N_e_* is fairly intuitive and will react to biological phenomena in expected ways (e.g. under population structure (Whitlock and Barton, 1997) or non-random mating (Caballero and Hill, 1992)).

LP state that the variables they introduce are “easily interpretable”. Their terminology, however, is not completely consistent nor coherent with more traditional uses, which can cause confusion. Given that *r* = *λ − μ* one might intuitively think that *r_p_* = *λ_p_ − μ_p_* but this is not the case. Pulled rates are simply different ways of summarizing congruence classes and each one is calculated using both speciation and extinction rates. *λ_p_* is reasonably intuitive, though given that extinction is also included it is more similar to a diversification rate than a speciation rate. Indeed, *λ_p_* is defined as the slope of the LTT plot (Louca et al., 2018) (see Fig. 3f, 4f), which corresponds to the past diversification rate, and to the speciation rate at present in the case all extant species are included (Nee et al., 1994).

Pulled diversification rate, however, is much more difficult to interpret, perhaps initially because the “pull” of *r_p_* is not the same as the “pull” of *λ_p_*. Whereas *λ_p_* decreases in value relative to *λ*, *r_p_* is delayed in time relative to *r* (Fig. 5) and could better be termed as “delayed” rather than “pulled”. We simulated a variety of diversification scenarios from simple to more complex (Fig. 5) and show that *r_p_* and *r* are similar in each case. However, *r_p_* is not as intuitive as *r* or *N_e_*. For example, drastic increases in *r* can lead to sharp decreases in *r_p_* (Fig. 5a). The inverted pattern *r_p_* presents in this case would make it challenging to present in a clear and concise way. Given the added difficulty of its interpretation we question whether *r_p_* provides a more useful estimate of the process of diversification than an estimate of *r*.

However, compared to other pulled rates, *r_p_* could be particularly useful, not as an effective parameter, like *N_e_*, but as a reasonable approximation of the true *r*. Indeed, we noted above that when shifts in *λ* are not too severe nor too rapid, *r_p_* is close to *r* (Fig. 5). Trying to biologically interpret fine-scale variations in *r_p_* would certainly lead to spurious conclusions. However, changes in *r_p_* at a large scale are good proxies for large scale variation in *r*. This is clearly illustrated in figure 5a where the main trend of the *r_p_* is a recent increase in diversification, and in figure 5d where the main trend is the stability of diversification despite rapid, short-term oscillations.

Pulled rates can be estimated using only the shape of the LTT plot, without any further information since they are non-parametric estimates that do not suffer from the unidentifiability problems outlined previously. However, they cannot be directly interpreted in biologically meaningful terms. To estimate rates that are meaningful (e.g. *λ*, *μ* and *r*), one needs to make further assumptions such as constant rates of speciation and extinction over time.

### On the use of models

The discussion sparked by Louca and Pennell (2020) highlights an important issue: evolutionary biologists should be interested in the actual history of diversification of the clades they study. The framework developed by Louca et al. (2018) shows how to do this using the shape of the LTT plot, without making strong assumptions about past speciation and extinction rates. Indeed, the slope and curvature of the LTT plot contain information about the diversification history of the clade. Much of the debate, however, focuses on the ability to recover a “true” history of diversification. Indeed, the goal of a scientific study should be to find out what really happened, but it becomes confusing if one considers a simulated birth-death process as the “true” history. This birth-death process is determined by two parameters (*λ* and *μ*) that can vary over time. These parameters are supposed to correspond to the rate that a lineage splits into two lineages, or goes extinct. In reality, however, a species does not have a speciation and an extinction rate in the same way it has a geographic distribution and a population size. These rates only make sense when they are aggregated over a number of species and a certain amount of evolutionary time. That is, they are descriptive statistics summarizing much more complex processes that are acting at the microevolutionary level, and that would eventually lead to speciation or extinction. Louca and Pennell (2020) convincingly show that one cannot estimate these statistics reliably from LTT plots, and propose statistics that can be estimated more reliably. That these alternative statistics do not exactly correspond to the parameters of the birth-death process is not a problem; the birth-death process is only a model of diversification, and not the truth about diversification itself. The framework built by Louca et al. (2018) and LP allows us to use the LTT to test whether the diversification rate was constant or not. If researchers want to know how speciation and extinction actually changed to give rise to this diversification history, they will have to use other methods.

## Conclusion

Louca and Pennell (2020) have pointed out key issues with how we approach macroevolutionary modelling, namely the inability to distinguish historical diversification scenarios under certain circumstances. Their formalization of the unidentifiability issues in LTT-based models is an important step forward that provides the mathematical tools to study the associated issues further. LP highlight the avenues we must consider and develop upon to ensure we do not make similar mistakes in the future. Whether variation in diversification rate is due to changes in speciation or extinction is certainly an interesting avenue of research, but LP have shown that exploring this would require much more than just fitting a model with speciation and extinction rates to an LTT. Indeed, more recent diversification models go beyond this by making use of additional information that is ignored by models relying only on the LTT. Awareness and consideration of potential unidentifiability issues is important for macroevolutionary biologists going forward when they employ such models of diversification. However, it is important to note that LP does not show that speciation and extinction cannot be estimated with evolutionary trees (Pagel, 2020). Instead, they show that when using extant timetrees with a single LTT-based approach, unidentifiability issues are encountered in the estimation of speciation and extinction rates, and that these problems can be circumvented by making use of pulled rates. Further work is needed to identify the extent to which the issues raised in LP apply to the more complex models of diversification used today. Comparisons should be made in empirical studies that use both traditional and pulled rates, to see if differences in results exist between these approaches in practice. In the meantime it is important that the field continues to grow by using and building upon modern macroevolutionary methods, albeit with a critical eye.

## Acknowledgements

This article was initially conceived from discussions held with the Macroevolution in Montpellier (MiM) group. We thank Stilianos Louca and Matt Pennell as well as Hélène Morlon and her research group for comments on an earlier version of this manuscript. We also thank Marcos Mendez, Sally Otto, Dan Schoen, Jürg Schönenberger and the rest of the DiveRS group for comments and discussion throughout. N.M. is supported by an “Investissements d’Avenir” grant managed by the “Agence Nationale de la Recherche” (CEBA, ref. ANR-10-LABX-25-01). T.L.P.C. is supported by funding from the European Research Council (ERC) under the European Union’s Horizon 2020 research and innovation programme (grant agreement No. 865787). E.L.R. and F.L.C. are supported by funding from the European Research Council (ERC) under the European Union’s Horizon 2020 research and innovation programme (project GAIA, agreement No. 851188).

## Supplementary Material

Code associated with this manuscript is available from

http:/github.com/ajhelmstetter/pulled_rates

## Notes

http:/github.com/ajhelmstetter/pulled_rates

### Competing Interest Statement

The authors have declared no competing interest.

